# Predicting individual skill learning, a cautionary tale

**DOI:** 10.1101/2022.04.24.489296

**Authors:** Dekel Abeles, Jasmine Hertzage, Moni Shahar, Nitzan Censor

## Abstract

People show vast variability in skill learning. What determines a person’s individual learning ability? In this study we explored the possibility to predict participants’ future learning, based on their behavior during initial skill acquisition. We recruited a large online multi-session sample of participants performing a sequential tapping skill learning task. We trained machine learning models to predict future skill learning from raw data acquired during initial skill acquisition, and from engineered features calculated from the raw data. While the models did not explain learning, strong correlations were observed between initial and final performance. In addition, the results suggest that in correspondence with other empirical fields testing human behavior, canonical experimental tasks developed and selected to detect average effects may constrain insights regarding individual variability, relevant for real-life scenarios. Overall, implementing machine learning tools on large-scale data sets may provide a powerful approach towards revealing what differentiates between high and low innate learning abilities, paving the way for learning optimization techniques which may generalize beyond motor skill learning to broad learning abilities.

## Introduction

People vary substantively in their ability to execute daily skills. What are the sources of such variability? Most studies have focused on initial and online task performance, known to vary between individuals (Anderson, Lohse, Lopes, & Williams, 2021). Thus, with no prior practice, some individuals might exhibit outstanding performance, while others might express slow and inaccurate performance. Importantly, people vary greatly in their ability to learn new skills as well, with the range of possible improvement differing between individuals. Predicting learning based on early skill acquisition offers an abundance of benefits and may be useful for effective adjustment of training regimes in daily life and for neurorehabilitation. What determines individual differences in learning abilities? Here, we aimed to investigate individual differences in skill learning by predicting the amount of learning an individual will exhibit across different time intervals, based on information extracted from performance at an early session.

Investigating individual differences with complex statistical modeling requires a large pool of participants. Therefore to address this question, we leveraged online platforms enabling crowdsourced recruitment producing large-scale data sets (Chandler & Shapiro, 2016; Ranard et al., 2014). Furthermore, the combination of such online platforms along the recent rise of machine learning models as means to understand rich data sets in neuroscience (Richards et al., 2019), provides a unique opportunity to investigate individual differences in skill learning.

To predict the extent of learning from skill acquisition characteristics, we utilized a common motor sequence learning task, widely used to model human skill acquisition (Brown & Robertson, 2007; Cohen, Pascual-Leone, Press, & Robertson, 2005; Genzel et al., 2012; Karni et al., 1998; Muellbacher et al., 2002; Perez et al., 2007; Reis et al., 2009; Robertson, Pascual-Leone, & Press, 2004; Wiestler & Diedrichsen, 2013; Wu, Srinivasan, Kaur, & Cramer, 2014).

Thus, we conducted a large-scale crowdsourced experiment, recruiting online participants to take part in 3 learning sessions, with a retention session following one week, and an additional long-term retention session following 2-4 months. First, we validated that online participation demonstrates common learning rates within each session as well as between sessions offline gains (Karni et al., 1995; Lugassy, Herszage, Pilo, Brosh, & Censor, 2018; Robertson et al., 2004). Next, we applied a wide array of machine learning models based on engineered features derived from existing literature of motor skill learning, as well as models based on raw data, using machine extracted features with no involvement of prior knowledge.

## Methods

### Participants

Participants were recruited online from the Amazon Mechanical Turk platform (https://www.mturk.com). Qualifications for registered MTurk workers to participate in the first session of the experiment were: above 95% approval rate in previous MTurk assignments, currently located in the USA, right-handed, and did not previously participate in a sequential tapping task from our lab. Each of the following sessions were made available to qualified participants according to the predefined scheduling scheme and was available for 12 hours. Data were collected using non overlapping batches of participants – session 1 of the experiment was made available on a Monday and the next sessions accordingly. This resulted in the following number of participants per session: Session 1: 571 participants, Session 2: 334, Session 3: 273, Session 4: 195, Session 5: 103. Additional exclusion criteria were enforced to make sure the remaining sample of participants were all attentive and complied with instructions (see below). This resulted in the final sample of: session 1: N=460; 274 Female; Mean age = 43.35, Std = 12.99; session 2: N=254; 154 Female; Mean age = 43.29, Std = 12.83; session 3: N=203; 116 Female; Mean age = 44.07, Std = 12.72; session 4: N=134; 75 Female; Mean age = 46.08, Std = 13.00; session 5: N=75; 39 Female; Mean age = 47.48, Std = 12.47. All participants used a button press to sign an online informed consent form presented at the beginning of each session. The payment scheme for all sessions was visible in the experiment page on the Mturk platform. To minimize dropouts, the compensation increased as sessions progressed (1.5$, 2$, 2.5$, 2$ for the shorter 4^th^ Retention session, and 5$ for the final long-term Retention session).

### Task

Participants performed a procedural motor task - the sequence tapping task (Karni et al., 1995), a highly common task used in numerous motor learning studies (Albouy et al., 2012; Bönstrup, Iturrate, Hebart, Censor, & Cohen, 2020; Herszage, Sharon, & Censor, 2021; Rickard, Cai, Rieth, Jones, & Ard, 2008). Participants were instructed (using illustrative slides) to place their non-dominant left hand on their keyboard in a one-to-one correspondence between fingers and digit-numbers; pinky – #1, ring finger – #2, middle finger – #3, index finger – #4. They were instructed to repeatedly tap the requested pattern (4-1-3-2-4) as fast and as accurate as possible using their left hand for the entire trial duration (10 seconds). A 10 second count-down screen preceded each trial and served as a break. Feedback was provided in the form of dots, with each keypress adding an additional dot to the display, regardless of correctness. Except for the sequence itself, this was the only visible item on the screen during the trial. The experiment was programed in Psychopy (Peirce et al., 2019) and was hosted on Pavlovia servers (https://pavlovia.org/).

### Experimental procedure

Before the first session, participants reported their age, gender, education level, time of weekly engagement with musical instruments and time engaged in physical activities. Additionally, at the beginning of each session, participants were asked to report the duration and the quality of sleep on the night preceding that session. At the end of each session, as a simple attention check, participants were asked to report the hand they used to perform the task. The study initially comprised of 4 sessions - each consisting of 36 trials except for the Retention session (4th session) containing 9 trials. A fifth session, the long-term Retention session, was made available 2-4 months after the completion of the Retention session, and comprised of 36 trials, identical to the first 3 sessions (figure 1a).

**Figure 1:**
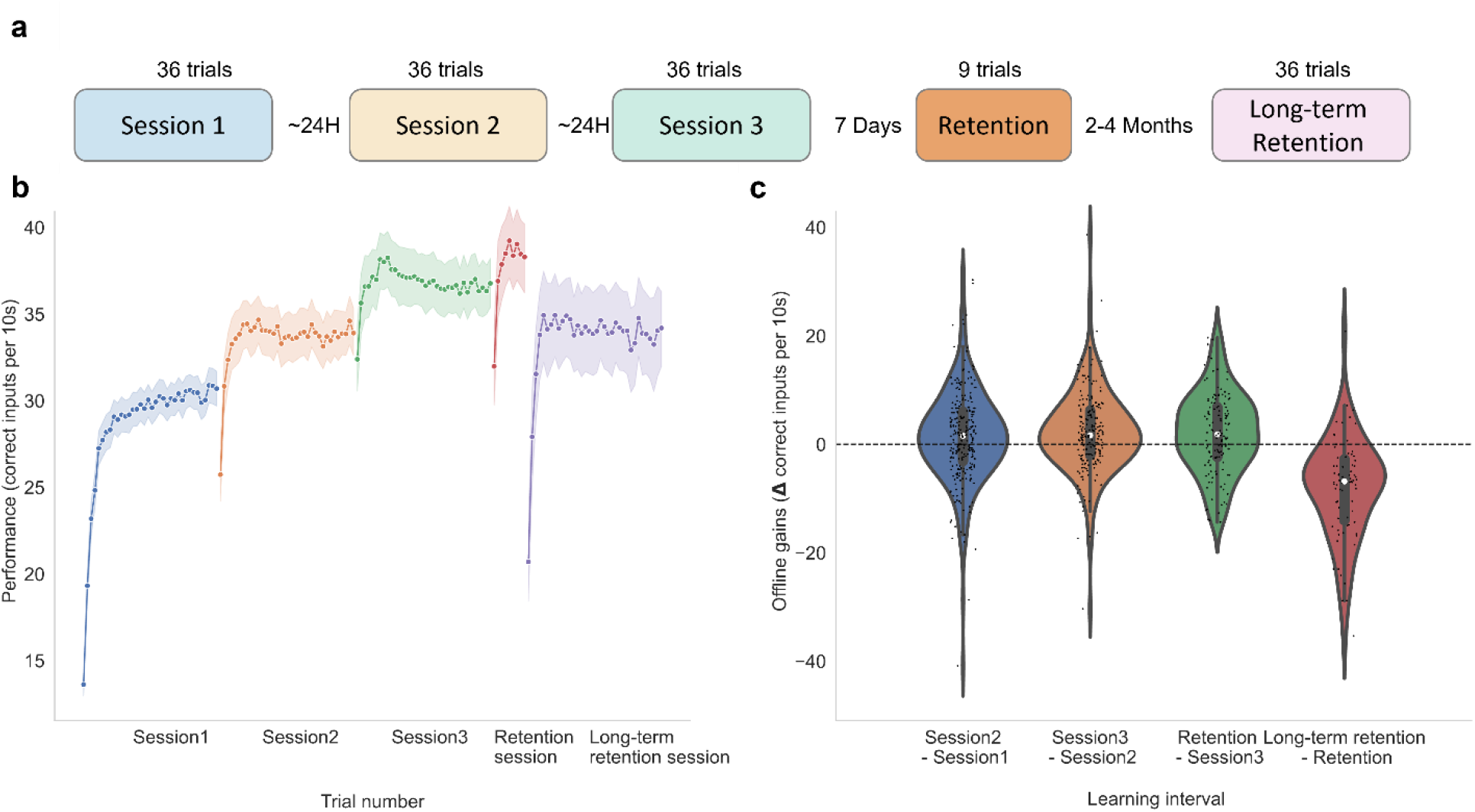
Task performance within and between sessions. a) Experimental design. b) Learning curves across all five sessions (session 1 – blue, session 2 – yellow, session 3 – green, Retention session – orange, Long Term Retention session – pink), the shaded area represents the 95% confidence interval. c) Offline gains between consecutive sessions. Data points in the violin plots represent offline gains for each participant. The white dot represents the median.

### Data analysis and machine learning feature engineering

All analyses were performed using custom code written in python (Van Rossum & Drake Jr, 1995). Data preprocessing and handling was done using the Numpy (Harris et al., 2020) and Pandas (McKinney, 2010) package. The machine learning pipeline was defined using Scikit-learn (Pedregosa et al., 2011) and Pytorch (Paszke et al., 2019). The Matplotlib (Hunter, 2007) and Seaborn (Waskom, 2021) libraries were used for data visualization. Statistical analysis was conducted using Pinguin (Vallat, 2018).

Participants were qualified to continue to the next session if they did not end the experiment mid-session and averaged at least 9 input characters per trial. Additionally, to validate participants’ attention to the task, data were discarded from all sessions if participants were too slow to start the trial following a break (first input exceeded 2 seconds) or failed to respond in more than 5 trials per session. Next, if the reported sleep duration was outside of the acceptable range of 6-12 hours, the data from that session and all following sessions were discarded.

Performance was defined as the overall number of correct keypresses in a trial (Censor, Horovitz, & Cohen, 2014; de Beukelaar, Woolley, & Wenderoth, 2014; Herszage et al., 2021; Korman et al., 2007). Keypresses were deemed correct if they were part of the complete requested pattern (4-1-3-2-4). If the trial ended mid-pattern, all keypresses from the start of that pattern were also considered correct. To minimize the effects of fatigue, *learning* was defined as the difference between the average of the 3 best trials in each session.

The following statistics were extracted from each session for each participant: *start performance* was defined as the average of trials number 2 and 3 (trial 1 not included due to warm-up decrements) (Adams, 1952; Rickard et al., 2008). *End performance* was defined as the mean of the last 3 trials in a session. *Maximal and minimal performance* were defined as the mean of the 3 trials with highest/lowest performance within each session. *Offline gains* were defined as the difference between consecutive sessions i.e., the *start performance* in session n+1 was deduced from the *end performance. Continuity* was defined as the average of the longest consecutive correct keypress of each trial across an entire session (Herszage et al., 2021). The *mean accuracy* was also computed for each participant in each session based on the average accuracies in all trials within the session. Additionally, the average response time of the first keypress of each trial across the session was defined as the *mean first RTs* and used as a proxy for estimating the level of attentiveness during the trial.

### Session dynamics

Session performance, defined as number of correct keypresses per trial within a session, was fitted with a learning curve according to the following equation:

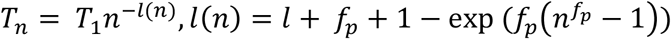

where *T* - the amount of correct keypresses, *l* - learning rate, *fp* - fatigue paramter (Asadayoobi, Jaber, & Taghipour, 2021), *n* - trial number. *Scipy*.*optimize*.*curve_fit* (initial guess for parameters (0.5,0.2,0) all bounded between [0-1]) was used to find the optimal Parameters *fp, l* and *T*_1_ for each participant and session.

### End of session slopes

A regression line (intercept and slope) was fitted for the number of correct trials for the last 15 trials in the session separately for each participant and session (excluding session 4, which included only 9 trials).

### Locally weighted scatterplot smoothing (lowess) features

For each participant, the correct number of keypresses per trial were smoothed across the session using a non-parametric local regression (*statsmodel*.*api*.*nonparamateric*.*lowess, fraq* = 0.5). Several features were extracted from the smoothed curve. First, we defined the regions of plateau on the curve as the longest streak of consecutive trials in which the derivative was below 0.25, meaning that the smoothed improvement between trials was less than a quarter of a keypress. The start and end of the plateau were defined as the first and last trials within this streak and the streak count was their difference. Additionally, the maximum of the smoothed curve and its index within the session (the trial in which it was achieved) were also extracted per participant and session.

### Within sequence consistency dynamics

To derive a representation of within sequence dynamics we first extracted the response time of the last sequence in a correct pattern (the 5^th^ input per sequence) in relation to the first input of the same sequence. This resulted in a vector of last keypress durations (locked to the first input of the sequence) for all correct sequences in the order of execution. To examine the consistency of this input over time we calculated the standard deviation over a running window of 10 consecutive inputs (*running RT consistency*). This running estimate was then fitted with a 3^rd^ degree polynomial (using the *numpy*.*polyfit* function). The coefficients of this polynomial and the fit prediction error (root mean square error) were used as additional hand-crafted features which capture the pattern dynamics across the session for each participant.

### Pattern consistency trend

To examine the amount of monotonicity apparent in the *running RT consistency* estimate, we used Spearman correlation with the corresponding vector of window number within the session. A high negative correlation suggests that a participant’s strategy gradually converged to a stable pattern. A high positive correlation on the other hand, suggests a diverged strategy, entering correct sequences less consistently as time progressed.

### Machine learning modeling

To test the predictive power of the behavior observed during initial training (session 1) on future learning induced by subsequent training sessions, three time intervals were examined: a) change in performance from the 1^st^ session to the 2^nd^ session. b) change in performance from the 1^st^ session to the 3^rd^ session. c) change in performance from the 1^st^ session to the 4^th^ retention session. Two additional time intervals were used to predict skill retention a) one week retention interval (from the 3^rd^ session to the 4^th^) and b) a long-term retention interval (2-4 months) (from the 4^th^ session to the 5^th^). Note that the number of participants decreases as the experiment reached later sessions, hence the number of observations available for modeling of later intervals is smaller. Accordingly, different modeling approaches were used, as detailed below.

The first approach utilized the engineered features as predictors and examined a wide range of machine learning techniques. Specifically, we tested: two tree-based models: Random Forest regression (Ho, 1995) and Sequential Regression Trees using gradient boosting (Xgboost) (J. H. Friedman, 2001). Regularized regression (Elastic net (Zou & Hastie, 2005)) and a multi-layer perceptron (MLP (Haykin, 1994)). Due to the large number of potential predictors, and to avoid over-fitting of the training set, we tested these pipelines both with and without an additional preprocessing step of principle components analysis (PCA)-based dimensionality reduction. Each modeling pipeline started with a standard scaler, transforming the feature values into z-scores. We used grid search for hyper-parameters tuning of the algorithms and regularization parameters. Each set of hyper-parameters was optimized separately for each type of algorithm, predictors step and time interval. The best model was selected based on the average 5-fold cross validation (CV) score. For each model type and time interval, the model selection was done in stages. In each stage an additional set of predictors was introduced based on their complexity, starting with high level features (i.e., session dynamic parameters) and ending with the simplest features (performance per trial). Initially, only non-behavioral features were included (i.e., Age and Gender). Next, predictors were introduced in steps. In the 1^st^ step parameters from the learning curve were introduced. The 2^nd^ step included the parameters extracted to capture *Within sequence consistency dynamics* and the *pattern consistency trend*. The 3^rd^ step included *Lowess based features*. The 4^th^ step included *session statistics*. The 5^th^ step included the micro-offline and micro-online features of the first 5 trials (Bönstrup et al., 2019). And the 6^th^ and final step, included the performance per trial for all trials in the session. For prediction purposes, normalization was done using the means and standard deviations of the variables in the training set. Additionally, we tested a recurrent Long Short-Term Memory (LSTM) network architecture in which the input was the most common end-point measure (de Beukelaar et al., 2014; Herszage & Censor, 2017; Herszage et al., 2021; Karni et al., 1995) of the task - the number of correct keypresses for each trial in the first session.

The second approach examined the prediction of future learning, based on all previous sessions. We used a linear regression model with correlation-based feature selection, introducing all available predictors at once and running a hyperparameters grid search on the number of selected features.

In the third approach, models were trained directly on raw data from the first session, predicting learning between the first and second training sessions. Task performance was represented as a binary image of size 4 × 7200, where rows represent the key identity (1-4) and columns represent the time where the key was pressed (in 50ms bins). For example, a key press on the key “3” performed 250ms after trial start, will have a value of 1 in the coordinate (3,5).

We then trained a convolutional neural network to predict learning. Hyper parameters of the topology and the optimization parameters were tuned manually. Similarly, a convolution encoder-decoder based method was built using the above binary session image as input, geared to reproduce the same image with a compact embedding layer which is then used as features in a regression analysis.

### Model evaluation

The parameters that resulted in the best performance on the training-set for each model type and prediction interval were used to re-train the model on the entire training set and examine it on the 20% of hold-out data that was not accessible during training. The final score is thus the reported explained variance (R^2^) of the hold-out dataset.

### Statistical analysis

One sample t-tests were used to examine the statistical significance of the offline gains analysis. Correlational analyses were conducted using Pearson or Spearman correlation.

## Results

We first validated that performance was consistent with previous studies employing the same task in laboratory settings (de Beukelaar et al., 2014; Herszage & Censor, 2017; Karni et al., 1998; Korman et al., 2007). Indeed, participants displayed typical learning curves (figure 1b), with significant learning expressed both within-session, and between-sessions as offline gains (Karni et al., 1998; Press, Casement, Pascual-Leone, & Robertson, 2005; Walker, Brakefield, Morgan, Hobson, & Stickgold, 2002) (figure 1c). Specifically, there were significant offline gains between sessions 1 and 2 (*t*(253) = 2.639, *p* = 0.009, *Cohen’s d* = 0.126, *CI* = [0.36 2.45]), and between sessions 2 and 3 (*t*(202) = 4.008, *p* < 0.001, *Cohen’s d* = 0.191, *CI* = [1.08 3.16]). Interestingly, even when the skill memory was tested following one week, additional offline gains were evident, with a significant improvement between session 3 and Retention session 4 (*t*(133) = 3.154, *p* = 0.002, *Cohen’s d* = 0.183, *CI* = [0.75 3.28]). In addition, during the long term retention interval, lasting between 2-4 months (see *Methods*) a significant reduction in performance was observed (difference from Retention (4^th^ session) to Long-term Retention (5^th^ session): *t*(74) = -7.661, *p* < 0.001, *Cohen’s d* = 0.722, *CI*= [-10.32 -6.06]), indicating a decay of the memory trace over a period of months. Overall, these results validate typical within and between session motor skill learning.

How could machine learning tools be applied to predict future learning? We first used ML with engineered features (see *Methods*), training discriminative algorithms to predict learning based on performance in the first session. To that end, our goal was to predict the improvements between performance in session 1 and performance in each of the subsequent sessions 2-4. To minimize within session effects of warm-up and fatigue (Adams, 1952; Rickard et al., 2008), between–session learning was quantified based on maximal performance in each session (see *Methods*). Potential predictors were introduced in steps with diminishing feature complexity, ranging from whole session dynamics descriptors, to the number of correct keypresses in each trial. The best performing model was selected based on its mean cross validation and tested on a predetermined hold-out set. Models did not predict learning in the hold-out set (*session2 - session1*: *R*^*2*^ = 0.08, *R*^*2*^ =0.15; *session3 - session1*: *R*^*2*^ = 0.09, *R*^*2*^ *=-0*.*18; Retention session 4 - session1: R*^*2*^ = 0.01, *R*^*2*^ = 0.07) (Figure 2a). Of note, a negative R^2^ score indicates that model predictions do not explain any variance in the dependent variable.

**Figure 2:**
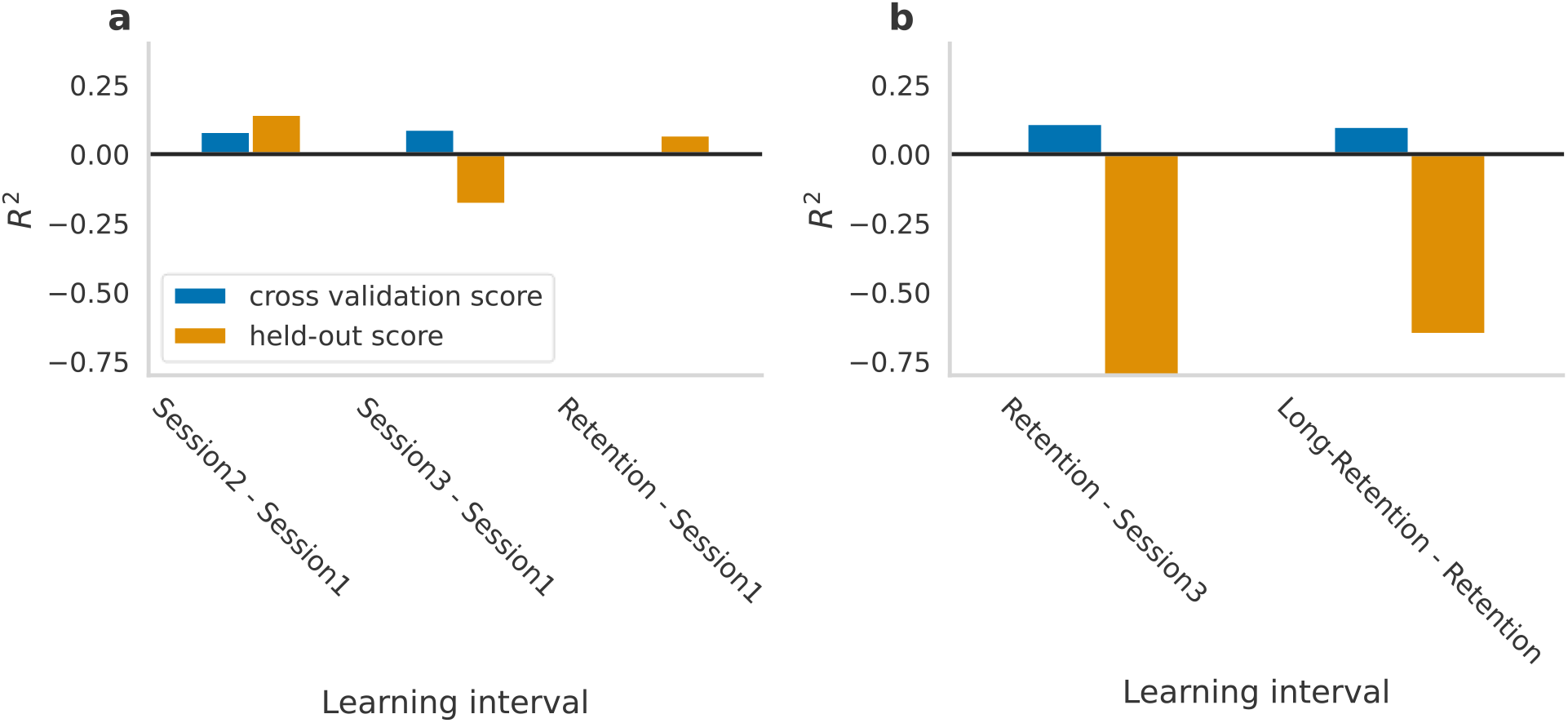
Model performance with engineered features. a) maximum mean cross-validation R^2^ scores (blue) and the corresponding hold-out R^2^ scores (orange) for each learning interval (X axis). b) Maximum mean cross-validation R^2^ scores (blue) and the corresponding hold out R^2^ (orange) for the two retention intervals (x axis).

Is behavior at initial stages of skill acquisition indicative of skill retention? To address this question, models were trained to predict the performance change during the short (from session 3 to Retention session) and long retention intervals (from Retention to Long-term retention), based on performance in either the first or all 3 prior sessions. The change in performance over both retention intervals was not predicted by the best performing model (highest cross validation score) as reflected in the negative *R*^*2*^ in the hold-out set (*Retention session - session3*: *R*^*2*^_*mean_cv_score*_= 0.11, *R*^*2*^_*test*_ *=* -0.84; *Long-retention – Retention session: R*^*2*^_*mean_cv_score*_= 0.10 *R*^*2*^_*test*_ *=* -0.65, figure 2b). Since the long-term retention interval showed negative changes in performance, further investigation of the data revealed that maximum performance in the Retention session was the best predictor for the subsequent long-term retention interval (Pearson’s *r*(73) = -0.49, *p* < 0.001, *CI* = [-0.65,-0.30]). Considering that maximum performance in the Retention session reflects both innate abilities and the overall benefit of training throughout the experiment, we examined the correlation between total learning and retention. Pearson correlation confirmed that the amount of total learning throughout the experiment (performance differences between session 1 and Retention session 4) was even a stronger predictor of the change in performance (Pearson’s *r*(73) = -0.58, p<0.001, *CI* = [-0.71,-0.40]), suggesting that participants exhibit long-term decay of their own learning before the retention interval.

Next, we tested whether a different approach of machine learning models, avoiding feature selection based on prior assumptions, will achieve better prediction of future learning. To further investigate prediction in that direction, we trained a convolutional neuronal network on data from session 1, represented as a binary matrix of size 4 × 7200, where rows represent key identity and columns represent keypress time within the session in 50ms time bins (Figure 3).

**Figure 3.**
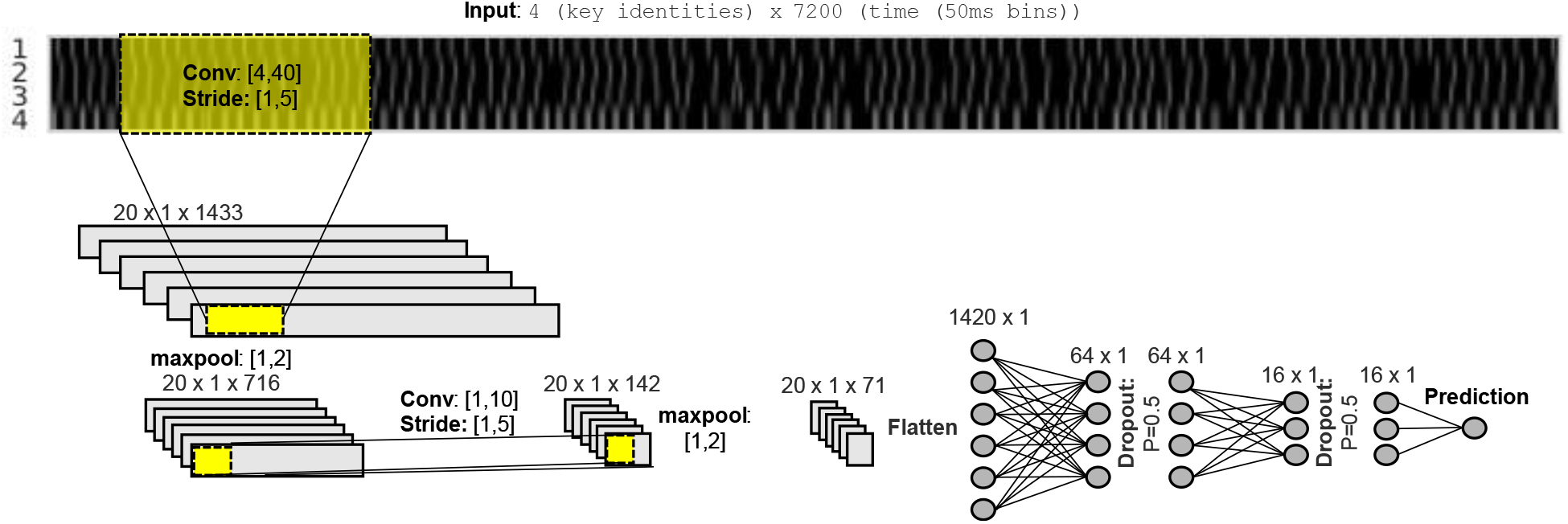
Convolution based neural network architecture. Input was represented as a 4 × 7200 binary matrix, where rows represent key identity (1-4) and columns represent time within the session (in 50ms time bins). The network architecture consists of two convolution layers, each followed by a pooling operation which is followed by 3 fully connected layers. The Rectified linear unit (Relu) was the selected activation function.

This representation reflects the available raw data, without imposing any definition of key correctness. This analysis was focused on the prediction of learning between the first and the second session, which includes the largest pool of participants. Additionally, to better utilize all available data, evaluation of model performance was based solely on cross validation. The best model resulted in mean cross validation *R*^*2*^_*test*_=-0.049, std = 0.053 performance. Consistent with this result, two additional models, using a convolution based encoder-decoder and LSTM architectures (see *Methods*), did not show predictive power.

To further investigate the above results, we assessed the consistency of simple performance metrics in each session and between-session learning, using Pearson correlations. Performance in each session explained a large portion of the variance in Performance scores across the 3 sessions and Retention session (R^2^ range = [0.25-0.91], all p < 0.001; see figure 4a), indicating high test-retest reliability and thus a stable measure of individual performance. However, performance hardly explained any portion of the variance in learning (R^2^ range = [0.00,0.05]; figure 4b). While these results suggest that variability in performance can be explained by performance in previous sessions, variability in learning can hardly be explained. To further illustrate this point, participants were separated into 5 quantile ranges (each spanning 20%)(Stafford & Dewar, 2014) based on their maximum performance in the Retention session, plotted throughout the experiment (figure 4c). The plotted curves show that participant’s relative performance remained stable throughout the experiment.

**Figure 4:**
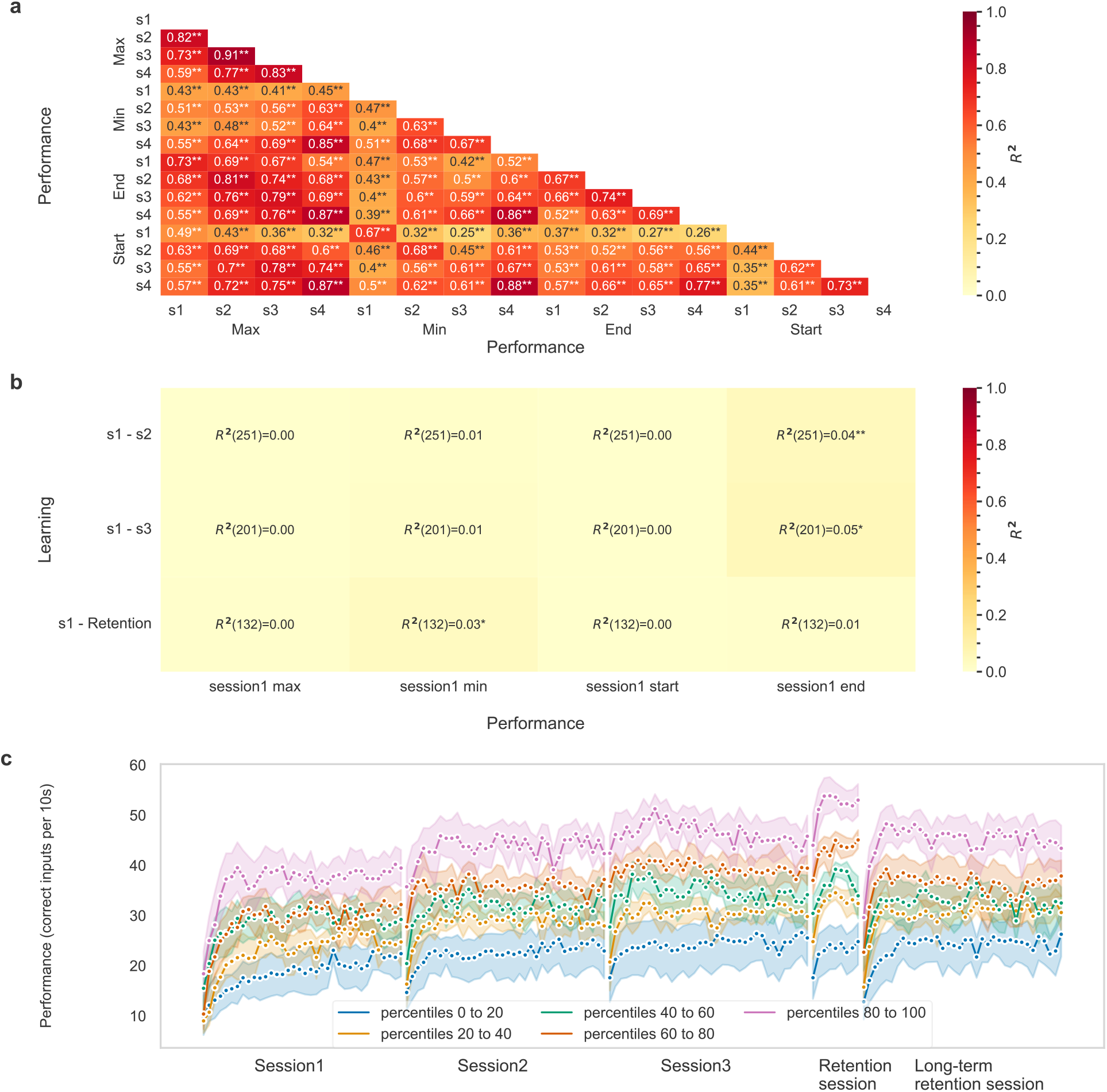
Performance was consistent across sessions but did not predict learning. a) Performance in all sessions explains a large portion of the variability in performance (*R*^*2*^ range = [0.25, 0.91]. b) Performance hardly explains the variability in learning (*R*^*2*^ range = [0, 0.05]. c) Performance throughout the experiment separated according to the performance quantile in the Retention session (colors), showing that participants’ relative performance rank remains stable across sessions. Shaded areas represent the 95% confidence interval. Statistical significance is marked with * for p<0.05 and with ** for p<0.001

## Discussion

The goal of this study was to identify what determines an individual’s skill learning ability, based on their initial behavior during skill acquisition. Learning was measured at different intervals, using large-scale crowdsourced data. Results showed that performance in early sessions did not predict subsequent learning, while variability in performance was explained by performance in previous sessions. In addition, participants exhibited long-term skill memory decay, bound by their own learning before the retention interval.

Machine learning techniques were leveraged to predict learning, utilizing several families of algorithms relying both on manually engineered features and on raw data representations. First, we extracted various features from the observed behavior in the task, ranging from high level features such as the parameters of the learning curve, to simple features such as the correct number of keypress in a trial. The applied models cover a wide array of approaches: Random Forest regression and Xgboost use an ensemble of weak learners and aggregate their predictions either based on consensus (random forest regression) or in a sequential manner. Multi-layered Perceptron (MLP), on the other hand, is a simple deep learning architecture consisting only of fully connected layers. The main advantage of these algorithms is their ability to capture interactions and other non-linear effects between predictors without explicitly modeling them by creating new variables. Two linear regression techniques were also examined due to their straightforward interpretability. Specifically, ElasticNet uses both L1 (Lasso) and L2 (Ridge) regularization penalties to limit model complexity while maintaining the linear relation between features and target. Finally, more sophisticated deep learning techniques were examined due to their ability to extract useful features from the data, without relying on expert knowledge and feature engineering.

A prerequisite of successful prediction of individual differences is a reliable test-retest metric for prediction (Spearman, 1961). This concept was demonstrated in other fields, such as the field of attentional control, where many canonical tasks, including Stroop (Stroop, 1935), Flanker (Eriksen & Eriksen, 1974), and Navon (Navon, 1977) result in robust between-conditions experimental effects, but in unreliable estimates of individual effects (Hedge, Powell, & Sumner, 2018), thus limiting insights regarding individual differences. Spearman and colleagues attributed this limitation to the calculation of a composite score as the difference between two measurements for the same individual (affecting test-retest reliability, Cronbach & Furby, 1970; Spearman, 1961). Critically, such differences between two measurements are the key outcome for evaluating skill learning. Therefore, while skill learning tasks have extensively shown robust and replicable results when examined between conditions (de Beukelaar et al., 2014; Gabitov et al., 2017; Herszage & Censor, 2017; Herszage et al., 2021; Korman et al., 2007), insights into individual differences may be limited. Accordingly, while large sample sizes may reduce standard errors and enable to detect average between-conditions effects, they do not necessarily improve the reliability of individual effects. This issue could be addressed by increasing the number of repeated measures or trials for each participant, as done for example in studies of perceptual learning (Sagi, 2011).

Furthermore, our analysis revealed that separating participants into 5 groups based on their performance in the Retention session, resulted in a visible, consistent classification throughout all sessions, suggesting that future learning may be too small to change participants’ rank.

Participants showing higher performance at the beginning, will also result in better performance at the end of the experiment. These results are consistent with previous findings of a large online sample of participants playing a complex online shooter game (Stafford & Dewar, 2014). When participants were split into 5 quantile ranges based on their best performance the curves remained separated from the very beginning of the task. Development of novel model motor skill tasks with high variability in between-session learning, and in which future performance is not determined by initial performance, may overcome the above constraints and provide further insights regarding learning variability, important for real-life scenarios. These may be combined with potentially useful predictors from other domains (Ackerman, 1987; Anderson et al., 2021; Chen, Gully, Whiteman, & Kilcullen, 2000), functional and anatomical neuroimaging information (Tomassini et al., 2011), or high-resolution kinematic inputs (Friedman & Korman, 2012).

In correspondence with other empirical fields testing human behavior, canonical experimental tasks developed and selected to detect average effects may constrain insights regarding individual variability, relevant for real-life scenarios. Accordingly, development of novel tasks with high test-retest reliability which model real-life learning, may shed light on the underlying mechanisms of individual differences in skill learning and promote personalized learning regimes geared to enhance human performance. Consequently, collecting large online datasets of behaving participants combined with advanced machine learning approaches, holds great potential for modeling future learning based on easily observable behavior during initial training. In turn, this may allow efficient resource allocation and enhancement of training regimes tailored to each person according to their innate abilities.

## Data and code availability

All collected data and the code for analysis are available upon request.

